# ARGfore: A multivariate framework for forecasting antibiotic resistance gene abundances using time-series metagenomic datasets

**DOI:** 10.1101/2025.01.13.632008

**Authors:** Joung Min Choi, Monjura Afrin Rumi, Peter J. Vikesland, Amy Pruden, Liqing Zhang

**Author notes:** Contributing authors. These authors contributed equally to this work (Joung Min Choi, Monjura Afrin Rumi).

## Abstract

**Background:** The global spread of antibiotic resistance presents a significant threat to human, animal, and plant health. Metagenomic sequencing is increasingly being utilized to profile antibiotic resistance genes (ARGs) in various environments, but presently a mechanism for predicting future trends in ARG occurrence patterns is lacking. Capability of forecasting ARG abundance trends could be extremely valuable towards informing policy and practice aimed at mitigating the evolution and spread of ARGs.

**Results:** Here we propose ARGfore, a multivariate forecasting model for predicting ARG abundances from time-series metagenomic data. ARGfore extracts features that capture inherent relationships among ARGs and is trained to recognize patterns in ARG trends and seasonality.

**Conclusion:** ARGfore outperformed standard time-series forecasting methods, such as N-HiTS, LSTM, and ARIMA, exhibiting the lowest mean absolute percentage error when applied to different wastewater datasets. Additionally, ARGfore demonstrated enhanced computational efficiency, making it a promising candidate for a variety of ARG surveillance applications. The rapid prediction of future trends can facilitate early detection and deployment of mitigation efforts if necessary. ARGfore is publicly available at https://github.com/joungmin-choi/ARGfore.

## 1 Background

The spread of antimicrobial resistance (AMR) is a global public health threat, undermining the efficacy and reliability of antibiotics to treat, cure, and prevent deadly bacterial infections worldwide [1, 2]. In 2019 and 2021, it was estimated that global deaths attributed to bacterial resistance to antimicrobials were 4.95 million and 4.71 million, respectively [3, 4]. Environmental dimensions of the antibiotic resistance problem are increasingly being recognized. Resistant bacteria and their antibiotic resistance genes (ARGs) can originate from natural environments, such as soil, and become amplified and mobilized as a result of human activities. Robust and coordinated surveillance systems targeting key nodes of evolution and dissemination of ARGs in the environment could prove critical in identifying new or re-emerging threats, predicting future risks, and implementing timely interventions [5]. Longitudinal studies across various ecosystems—including air [6], soil [7], sewage [8, 9], and wastewater [10–12]—have provided valuable insights into the temporal dynamics of ARGs and the factors contributing to their evolution, horizontal transfer, selection, and spread across ecosystem boundaries. Such understanding could prove crucial towards informing policy and practice aimed at optimizing antibiotic use and limiting the spread of resistance. A key challenge, however, lies in predicting the future proliferation of antibiotic resistance in the environment of interest. Such predictions, referred to as a “time-series forecasting problem,” are vital as resistance forecasting has the potential to enable timely responses to emerging outbreaks, allowing for containment strategies and more informed and targeted efforts to treat resistant infections in affected communities.

Recent studies have attempted to utilize statistical approaches to predict AMR using longitudinal data. For instance, Guo et al. [13] employed an Auto-Regressive Integrated Moving Average (ARIMA) model to forecast AMR among Gram-negative bacteria based on 15 years of data on antimicrobial consumption and resistance in a hospital setting. However, ARIMA based models assume a linear relationship between past observations and future predictions [14, 15] and thus may fail to capture complex non-linear patterns in antibiotic resistance that occur due to seasonal variation [16, 17]. Kim et al. [18] utilized Seasonal ARIMA (SARIMA) to predict the future burden of antimicrobial resistance (i.e., growth rate of resistance to their corresponding antimicrobials based on Minimum Inhibitory Concentration interpretations) for five bacterial pathogens prevalent in swine farms. While SARIMA aims to address seasonal variation, it is a univariate model. Forecasting ARG abundance, on the other hand, is a multifaceted issue due to the co-occurrence of ARGs in the environment. For example, ARG dissemination is thought to be driven in part by anthropogenic contamination of antibiotics and other antimicrobials (e.g., heavy metals) [19–21]. Such selective pressure can act directly on ARGs encoding resistance to corresponding antibiotic classes, or can indirectly co-select for ARGs encoding resistance to other antibiotics because they co-occur on the same genetic element [22, 23]. Horizontal gene transfer facilitated by mobile genetic elements (MGEs) can also result in the co-occurrence of multiple ARGs encoding resistance to multiple antibiotics [24, 25]. Such co-occurrences and co-mobilities among ARGs, resulting in both multivariate inputs and multivariate outputs, challenge forecasting efforts. Vector Autoregressive Moving-Average (VARMA), a variant of the ARIMA model, can handle multivariate data but is not equipped to handle seasonal variation. Additionally, for effective mitigation and pandemic prevention, approaches are needed that can anticipate threats well in advance. This calls for the implementation of a method using a multi-step-ahead approach, which could enable detection of early signs long before problems reach critical levels. While traditional time series models like ARIMA and its variants can provide some predictive power, they are likely insufficient for capturing the complex, seasonal, and multivariate nature of AMR data.

Recently, deep learning models have gained traction in solving forecasting problems that are multivariate and multi-output in nature [26–29]. Various neural network models that capture temporal dependencies between observations obtained at different time points have been developed for time-series forecasting. For example, recurrent neural networks (RNN) and its variants, such as Long Short-Term Memory (LSTM), Gated Recurrent Unit (GRU), or those combined with attention neural networks, have found application in diverse domains such as weather forecasting [30, 31], financial trend analysis [32, 33], power usage forecasting [34–36], and disease prediction [37, 38]. However, despite good performance, the computational demands of these RNN-based models result in low time efficiency due to the substantial costs associated with hyperparameter optimization that are required to capture long-range dependencies. In comparison, neural network models based on Multi-Layer Perceptron (MLP) stacks that implicitly capture input-output relationships outperform the latest RNN models while maintaining computational efficiency. Notably, N-BEATS introduced a deep neural network architecture with multiple stacks of fully connected layers that is adaptable to various domains, enabling rapid training [39]. Similarly, N-HiTS presented a comparable architecture with multiple MLP stacks, redefining stack structures by decomposing input signals into different frequencies through multi-rate data sampling with each stack predicting future signals at different rates and the outputs synthesized through multi-scale interpolation [40]. These RNN-free strategies have demonstrated significant improvements in accuracy and computational complexity for time-series forecasting. However, their application has predominantly been tested in univariate settings on benchmark datasets; such as weather and traffic, rather than in multivariate datasets with implicit interactions, such as biological datasets where many features exhibit inherent relationships.

Here we introduce ARGfore, a deep neural network-based forecasting method for predicting ARG abundances from time-series metagenomic data. From a multi-ARG input dataset, features incorporating inherent relationships among ARGs within the same drug classes are extracted using convolutional neural network (CNN) modules. These features are then fed into the forecasting module, consisting of three blocks with multiple MLP layers. The three blocks aim to decompose the input into the trend, seasonality, and residue components, learning the temporal patterns of ARG abundances in each component. The forecast outputs from the three blocks are aggregated to yield the final prediction of future ARG abundances. Compared to standard time-series forecasting methods, ARGfore demonstrated the lowest mean absolute percentage error across three distinct metagenomic datasets. In addition, ARGfore enhances computational efficiency of future ARG abundance prediction, making it suitable for real-time applications.

## 2 Methods

ARGfore consists of two modules: the drug class-based feature extraction module and the time-series forecasting module. The workflow is illustrated in Figure 1.

**Fig. 1.**
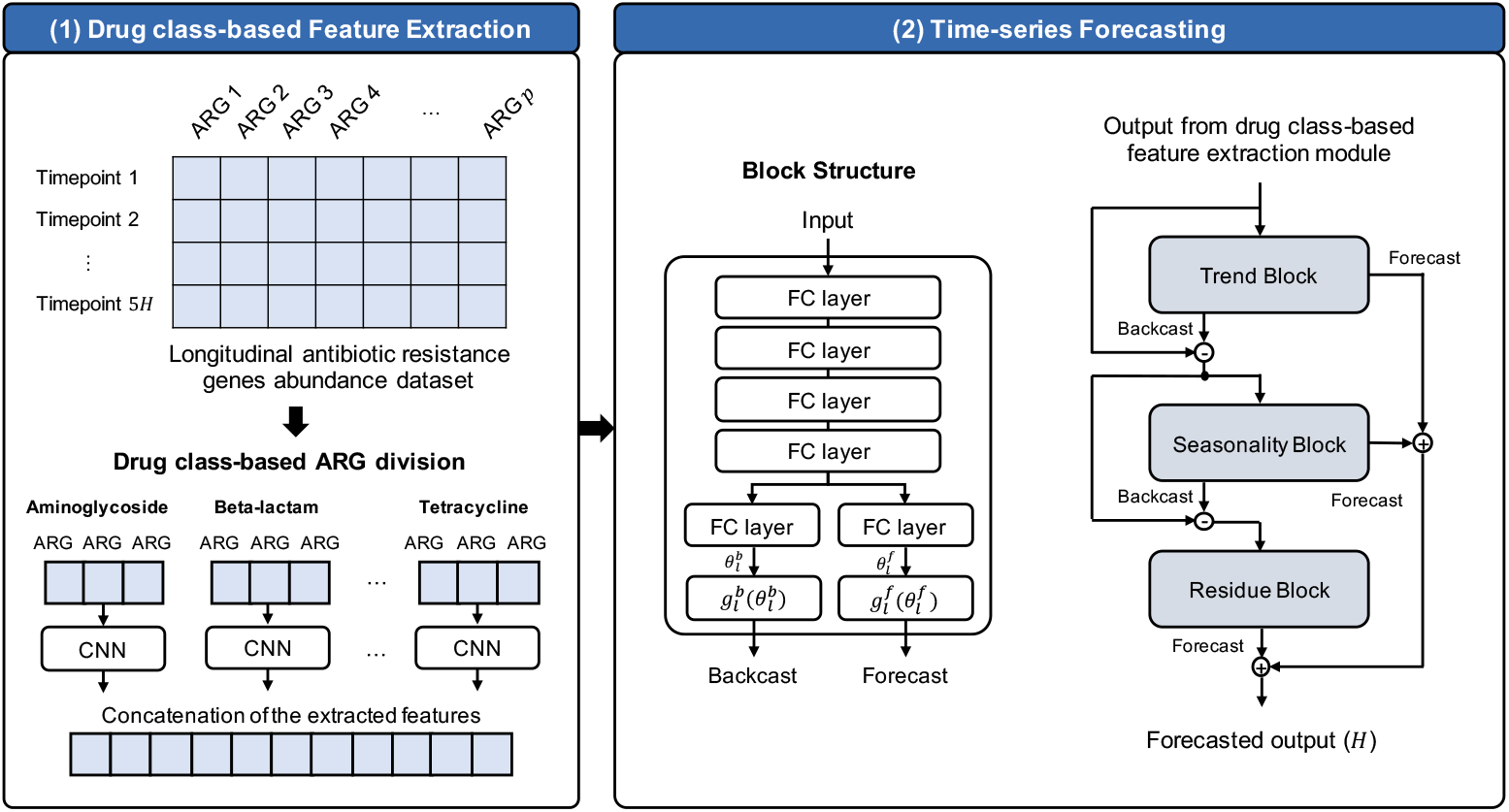
The ARGfore workflow for forecasting the abundance of ARGs using time-series ARG abundance data.

### 2.1 Drug class-based feature extraction

Co-occurrence of ARGs can stem from being carried by the same MGEs, encoding resistance to the same drug class [41, 42]. To capture such complexities, a CNN-based feature extraction module was introduced. In this module, ARGs from the input data were first grouped into clusters based on the drug classes to which they confer resistance. Spearman correlation matrix (*p × p* matrix for *p* ARGs in a cluster) was computed between ARGs for each cluster. Each row in the matrix was reduced to the geometric mean of correlation coefficients using the formula below (Equation 1).

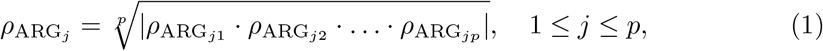

where *ρ*_ARG*jp*_ is the Spearman correlation between ARG_*j*_ and ARG_*p*_.

To establish similarity between neighboring ARGs, the ARGs within each drugclass-based cluster were sorted in decreasing order of the geometric mean of correlation coefficients. The CNN module was then applied individually to each cluster, extracting features. The module consisted of one 1D-CNN layer with 8 filters of size 3 by 3, followed by a Leaky ReLU activation function and a max-pooling layer. Extracted features from each cluster were concatenated and transferred to the trend block in the forecasting module.

### 2.2 Time-series forecasting

The time-series forecasting module was constructed based on the architecture of the N-BEATS model [39]. Let *H* be the length of time points for the forecast period that the model will predict. Recent papers [39, 40, 43] set the length of the input as the multiple of the forecast horizon *H*, ranging from 2*H* to 7*H*. Here, 5*H* was set as the default length of the input, which achieved the highest forecasting performance in our experiments shown in the Results section. The time-series data were decomposed into three blocks: the trend, seasonality, and residue blocks. Each block had four fully connected (FC) layers with ReLU nonlinearities, producing forward 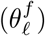 and backward 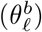 predictors of expansion coefficients as follows :

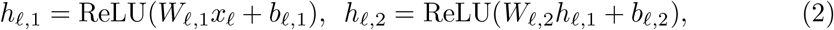

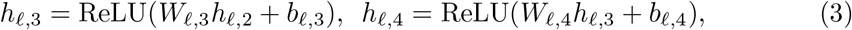

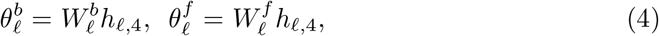

where *x*_ℓ_ is the input of the block ℓ, 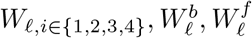, are the weights and *b*_ℓ,*i* ∈*{*1,2,3,4*}*_ are the bias terms for the block ℓ.

These coefficients were then transferred to the backward 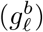 and forward basis layers 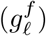 with pre-defined functions addressing the characteristics of each decomposition component, producing backcast output 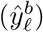 and forecast output 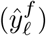 for the next *H* timepoints. The backcast output was excluded from the block’s input, and the resulting input was then fed into the next block. The forecast outputs from all blocks were aggregated hierarchically to provide the final forecast output.

#### 2.2.1 Trend block

The trend block, serving as the initial block and taking the output of the feature extraction module as input, is designed to capture the typical characteristics of trends in time-series data. Trends are often either monotonic functions or slowly varying functions. To adhere to this characteristic, the backward and forward basis layers 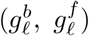 in the trend block were defined as a polynomial of a small degree *p*, representing a slowly varying function across the forecast window:

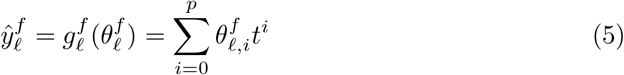

Here, 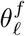 represents the polynomial coefficients predicted by the FC layers, *p* is set to 2, and the time vector *t* = [0, 1, 2, …, *H* − 2, *H* − 1]^*T*^ */H* is defined on a discrete grid ranging from 0 to (*H* − 1)*/H*, forecasting *H* steps ahead.

#### 2.2.2 Seasonality and residue blocks

To mimic the seasonal pattern inherent in time-series data – characterized by regular and cyclical fluctuations – the backward and forward basis layers 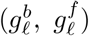 in the seasonality block were constrained to follow a Fourier series periodic function:

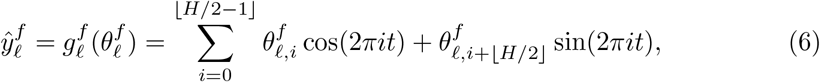

Where 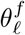 represents the Fourier coefficients predicted by the FC layers.

For the residue block, the backward and forward basis layers 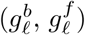 were defined simply as a FC layer :

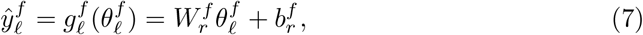

where 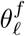 represents the outputs from the FC layers in the residue block, while 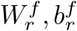 are the weight and bias terms, respectively. The forecast output from the trend, seasonality and residue blocks were collected and aggregated to provide the final forecast result.

Here, ARGfore was trained with the adaptive optimization algorithm Adam [44] with a learning rate of 10^−3^ based on the mean absolute error (MAE) loss for the actual and forecasted ARG abundance values. Model training ceased when the MAE loss did not decrease within 100 epochs. ARGfore was implemented using the Tensorflow library (Version 2.8.0).

### 2.3 Experimental design

#### 2.3.1 Data collection

To evaluate the performance of our proposed model, three longitudinal metagenomics datasets were utilized. The first dataset, referred to as ‘Influent’, consists of 98 samples collected from the influent sewage of a local 5 MGD wastewater treatment plant (WWTP) in Virginia, USA [46]. The second dataset, labeled ‘Effluent’, comprises 84 samples collected from the treated effluent of the same WWTP [46]. Both the Influent and Effluent samples were collected twice weekly between August 2020 and August 2021. The last dataset labeled as ‘Activated Sludge,’ which represents microbes enriched for their capability to break down organic pollutants in wastewater, was downloaded from NCBI [10]. It contains 98 activated sludge samples collected monthly from the aeration tank of the 300 MGD Sha Tin WWTP in Hong Kong, which treats saline sewage, spanning the period from June 2007 to December 2015. Nonpareil3 [45] was used to estimate library coverage for each sample using options -T kmer. The average coverage of Influent, Effluent, and Activated Sludge datasets are 73.26%, 73.94%, and 62.62% respectively.

#### 2.3.2 Preprocessing

A comprehensive pipeline for the microbiome dataset was designed to preprocess sequencing data, following the established standard metagenomics workflow [47, 48]. Raw read quality control was conducted using fastp [49], with parameters configured for (1) adapter trimming, (2) polyG tail trimming, (3) polyX tail trimming, (4) low complexity filtering, and (5) an average quality threshold ≥ 10. Subsequently, read decontamination was carried out using bbduk [50] and involved masking against human, rat, and cow genomes associated with the bbtools suite. Quantification of ARGs within the short reads was accomplished using DIAMOND [51], aligning them to the DeepARG database (DeepARG-DB) [52]. Calculation of 16S rRNA gene copies was performed by aligning short reads to GreenGenes v. 13.5 [53] with minimap2 [54]. The relative abundances of ARGs were determined as the ratio of functional gene copies to 16S rRNA gene copies [55], followed by min-max normalization. While we used 16S normalization in our data preprocessing, users have the flexibility to apply other normalization techniques as appropriate as long as the same normalization technique is used consistently for processing the data.

ARGs that were not present in all samples were filtered out in the Influent and Effluent datasets. In the Activated Sludge dataset, ARGs detected in at least 80% samples were kept as only a few ARGs were consistently present in every sample across the nine-year duration covered by this dataset. After preprocessing, 240 (Influent), 112 (Effluent), and 97 (Activated Sludge) ARGs remained. The typical sampling frequency of Influent and Effluent datasets was twice per week, resulting in a 3-4 day gap between samples. Some samples could not be collected on schedule due to logistical reasons, resulting in longer gaps (up to 14 days) in some cases. To mitigate the effect of missing values on model development, multivariate imputation by chained equations (MICE) was employed for the missing time points. To choose the most effective imputation strategy, we explored the impact of various imputation methods on ARG abundance prediction, comparing forecasting performance in the “Results” section. With imputation, a total of 112 (Influent), 107 (Effluent), and 102 (Activated Sludge) samples were used for our experiments.

#### 2.3.3 Clustering

The feature extraction module was tested by applying different clustering techniques. We deployed k-means clustering, hierarchical clustering, and clustering based on resistance to the same drug class or the same resistance mechanism. The information of the drug classes for input data was obtained from DeepARG-DB [52]. Resistance mechanism information was obtained from Comprehensive Antibiotic Resistance Database (CARD) [56] and CD-HIT [57] was used to cluster the ARGs in DeepARG-DB.

### 2.4 Comparison with other forecasting methods

We compared the performance of ARGfore to three categories of forecasting methods. First, we employed the state-of-the-art MLP-based forecasting model, N-HiTS. A stacked recurrent neural network-based model was also compared, comprising two LSTM layers with 64 and 32 units, followed by one fully connected (FC) layer. We also utilized the widely-used statistical approach for time-series forecasting, ARIMA. All the models are designed to work on multivariate datasets and for the statistical model we used multivariate variation of ARIMA namely VARMA. In the experiments, each dataset was split into an 8 : 2 ratio for training and testing, with models trained on the training dataset. The forecasting period *H* was set to 10 timepoints, and the input length for the model was set as 5*H* (50 timepoints). Performance evaluation metrics included the mean absolute percentage error (MAPE) and symmetric mean absolute percentage error (SMAPE), defined as follows:

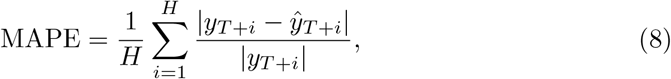

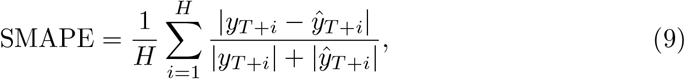

where *y*_*T*_ and 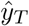 represent the actual and predicted value for time point *T*. In each experiment, MAPE and SMAPE were measured for testing datasets, with the normalized datasets scaled according to the original range of each feature value.

We also examined the impact of various imputation techniques during preprocessing step on ARG abundance forecasting and assessed the robustness of the models under different scenarios. Time-series imputation methods, including linear curve fitting, cubic curve fitting, and moving-window-based imputation (window size = 5) were implemented using Pandas library in Python. Additionally, we tested two widely-used imputation methods for handling missing data in longitudinal datasets, the Last Observation Carried Forward (LOCF) from imputeTS library [58] in R and MICE approach from scikit-learn.

## 3 Results

### 3.1 Performance evaluation of ARGfore

As summarized in Table 1, ARGfore consistently outperformed the other three forecasting methods (N-HiTS, LSTM, and ARIMA). ARGfore exhibited the lowest average MAPE of 0.308 and SMAPE of 0.306 when utilizing MICE imputation for the Influent dataset. LSTM demonstrated the second-best performance, with an average MAPE of 0.544 and SMAPE of 0.377, followed by N-HiTS and ARIMA. ARGfore also achieved the best forecasting performance for both Effluent and Activated Sludge datasets using MICE imputation, with an average MAPE of 0.772 and 1.075, respectively, compared to the second-lowest average MAPE of 1.116 and 1.204. Even when employing other imputation methods during preprocessing, ARGfore consistently exhibited superior forecasting performance compared to the alternative methods. These results suggest that our proposed model for forecasting ARG abundances in time-series metagenomic data is robust. Moreover, it was observed that all forecasting methods achieved the best performance in most cases when MICE imputation was employed. Therefore, we used MICE imputation for the subsequent experiments.

**Table 1.**
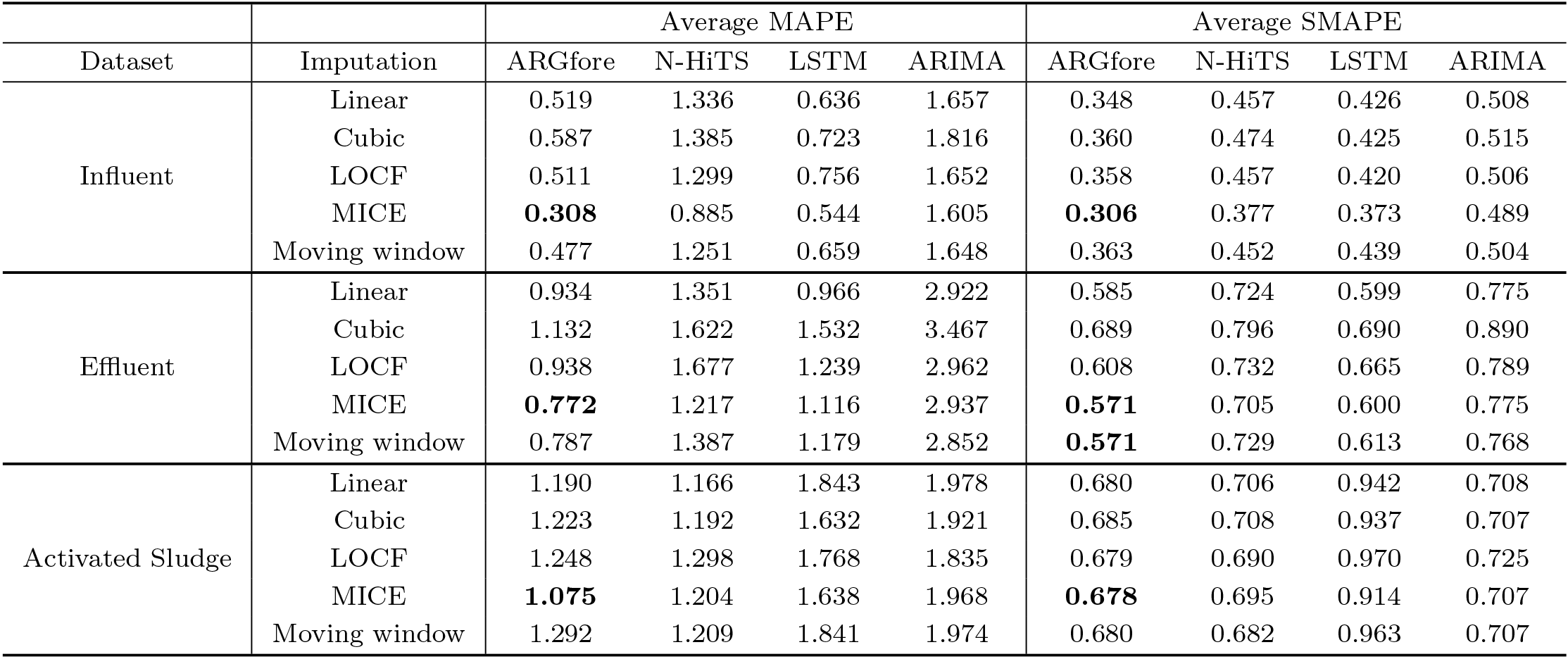
Forecasting performance results based on the average MAPE and SMAPE results using different imputation strategies for each of the three datasets.

We further examined the performance of these methods on individual genes. As an example, Figure 2 illustrates the forecasted patterns generated by various methods for the *tetA* gene in the Influent dataset. ARGfore demonstrated the closest alignment to the original pattern capturing the trends and fluctuations observed in the actual data (Pearson correlation coefficients between actual data and predicted by ARGfore, N-HiTS, LSTM, and ARIMA are 0.445, −0.112, −0.025, and 0.105, respectively). This highlights ARGfore’s superior performance in forecasting individual ARG dynamics over these methods.

**Fig. 2.**
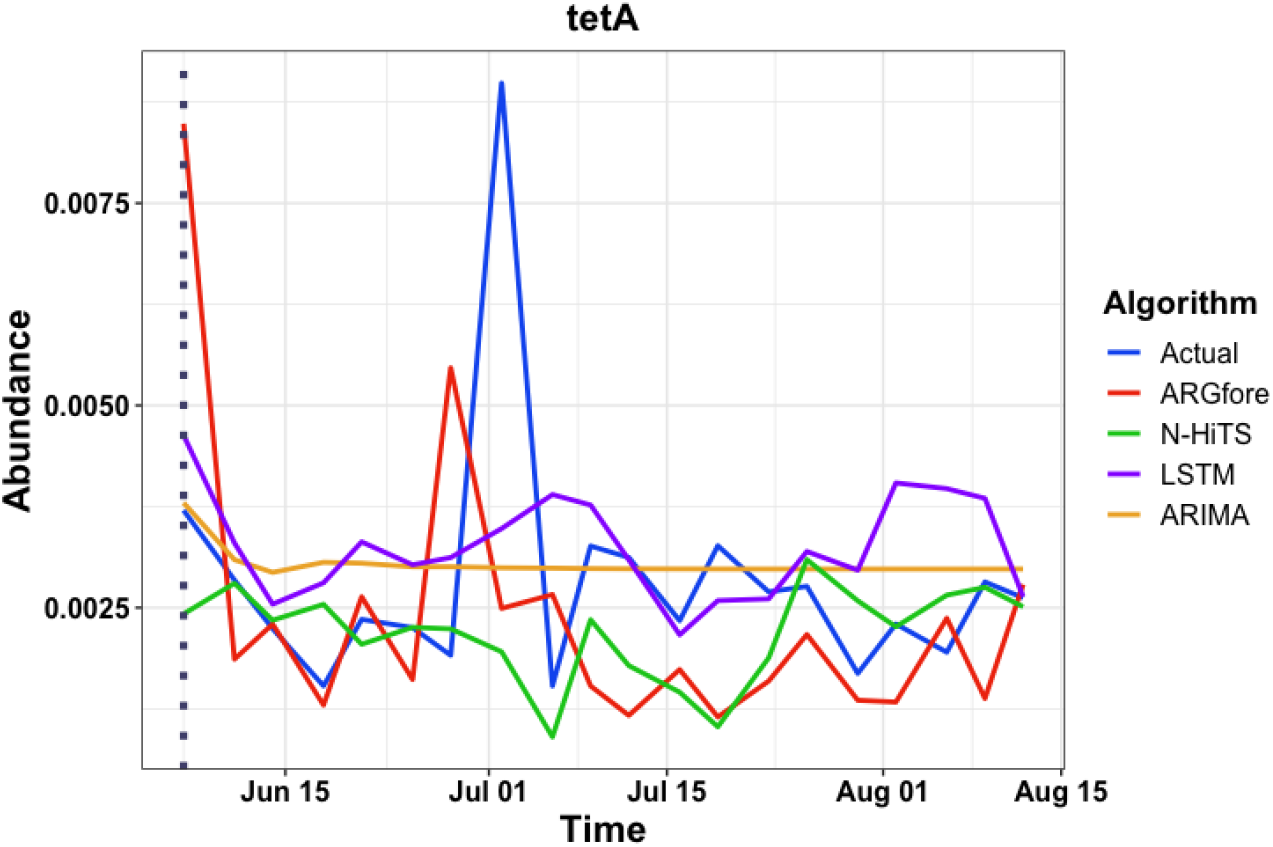
Forecasting *tetA* abundance in test data for Influent dataset using different time-series forecasting methods. Dataset was split into an 8:2 ratio for training and testing, with models trained on the training dataset. The trend predicted by ARGfore (red line) resulted in the strongest correlation with the actual trend (blue line) (Pearson, R=0.445).

During data pre-processing, we selected only those ARGs that were detected at all time points (i.e., 100% occurrence frequency) for both Influent and Effluent datasets, but 80% occurrence frequency for the Activated Sludge dataset. To examine the effect of this criterion on ARGfore performance, we selected ARGs for prediction based on three occurrence frequency requirements, 100%, ≥ 90%, and ≥ 80% presence. Table 2 shows that ARGfore is capable of predicting ARG sets with different occurrence frequencies; however, prediction accuracy increases when applied to ARGs with higher occurrence frequencies.

**Table 2.** Performance of ARGfore measured by the average MAPE and SMAPE for predicting ARG sets with different occurrence frequencies.

| Occurrence frequency | Average MAPE |  |  | Average SMAPE |  |  |
| --- | --- | --- | --- | --- | --- | --- |
|  | Influent | Effluent | Activated Sludge | Influent | Effluent | Activated Sludge |
| 100% occurrence<sup>a</sup> | 0.308 | 0.772 | 0.659 | 0.306 | 0.571 | 0.455 |
| 90% occurrence<sup>b</sup> | 0.615 | 1.268 | 1.012 | 0.385 | 0.617 | 0.63 |
| 80% occurrence<sup>c</sup> | 0.632 | 1.352 | 1.075 | 0.409 | 0.628 | 0.678 |

<sup>a</sup> 100% occurrence refers to ARGs detected at all time points, <sup>b</sup> 90% occurrence indicates ARGs detected in at least 90% of the samples, <sup>c</sup> 80% occurrence pertains to ARGs detected in at least 80% of the samples

### 3.2 Effectiveness of feature extraction in ARGfore

ARGfore employs an MLP-based model coupled with CNN for extracting ARG features relevant to drug classes to predict future ARG abundances. To assess the impact of feature extraction, we designed four variants of ARGfore by modifying or removing the feature extraction module from the model. The ‘w/o FE’ model eliminates the CNN-based feature extraction, while the ‘Drug class’ model utilizes drug class information for clustering. Similarly, the ‘Resistance class’ model uses resistance mechanism information to cluster the ARGs. The ‘K-means’ and ‘Hierarchical’ models perform K-means and hierarchical clustering, respectively, using the clustering results to group ARGs and extract features based on CNN. The Scikit-learn [59] Python package was employed for clustering, and the number of clusters (*K*) was determined using the Elbow method [60], resulting in values of 4, 3, and 2 for Effluent, Influent, and Activated Sludge datasets, respectively. Each dataset was divided into an 8 : 2 ratio for training and testing, and the experiment was repeated five times.

Figure 3 illustrates that, compared to the ‘w/o FE’ model, all four models with the feature extraction module achieved better forecasting performance. Moreover, the model using drug class information to group ARGs for feature extraction obtained the best performance in both Effluent and Influent datasets, with average MAPEs of 0.772 and 0.308, respectively. The ‘K-means’ model showed the best performance in the Activated Sludge dataset, with an average MAPE of 0.966, followed by the second-best ‘Drug class’ having the average MAPE of 1.076. These findings underscore the effectiveness of the drug class-based feature extraction method based on CNN in ARGfore, highlighting benefits for improving prediction of future ARG abundance values.

**Fig. 3.**
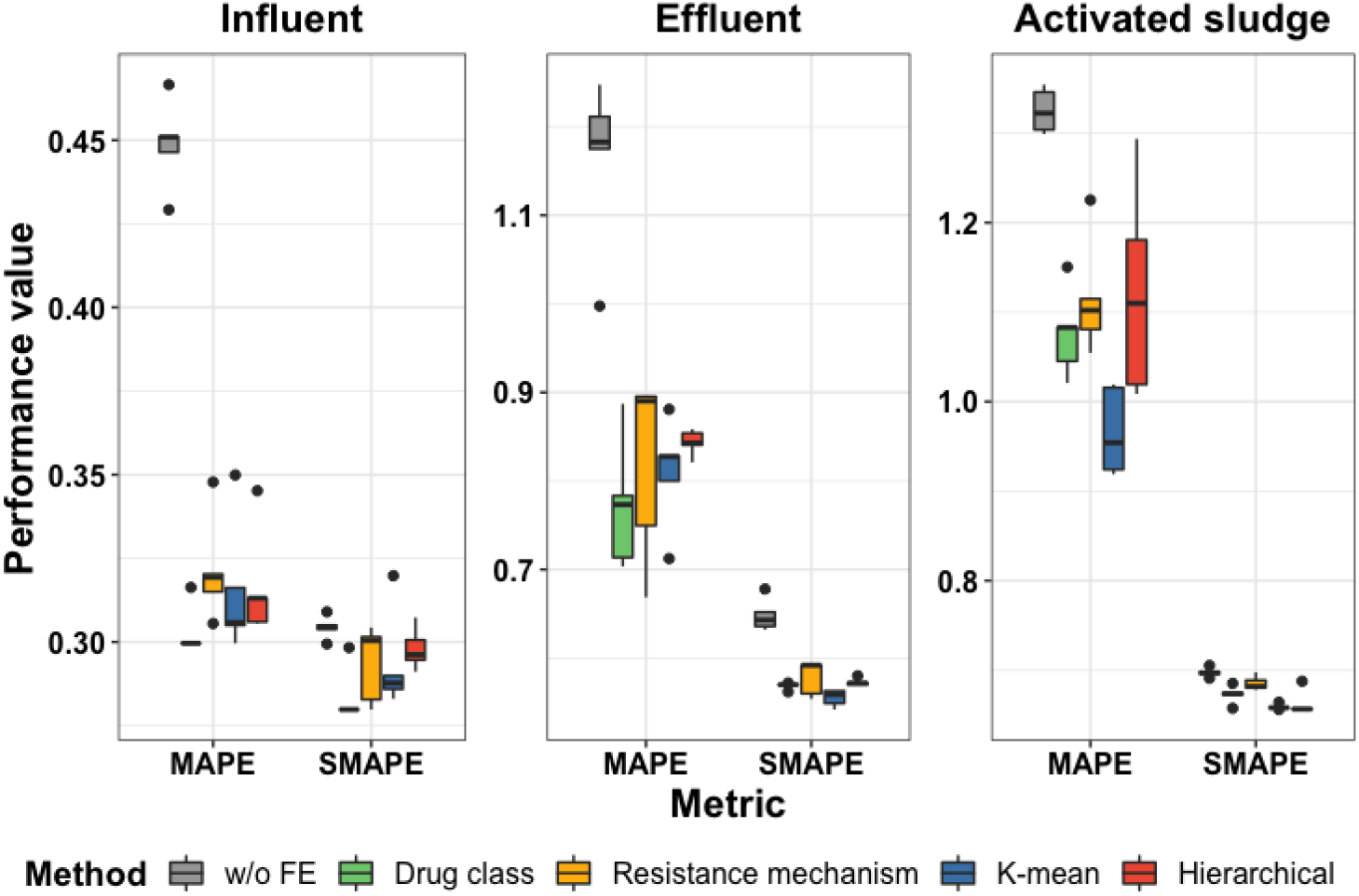
Time-series forecasting performance comparison of ARGfore and its variants with different clustering-based feature extraction methods using MAPE and SMAPE metrics. ‘w/o FE’ refers to without feature extraction, and ‘Drug class’, ‘Resistance class’, ‘K-means’ and ‘Hierarchical’ represents CNN-based feature extraction utilizing each clustering information, respectively. The lowest average MAPE and SMAPE was obtained for Influent and Effluent using drug-class based feature extraction method. For Activated Sludge dataset, k-means generated the lowest average and drug-class based clustering techniques generated the second lowest.

### 3.3 Run time comparison

We compared the run time of ARGfore with the other three forecasting methods. The average run time for each time-series dataset was measured by repeating the experiment five times. It was tested on a single-node server with 2 cores of Intel(R) Xeon(R) CPU @ 2.20GHz and 13 GB of main memory. Table 3 reveals that ARGfore exhibited the lowest average run time at 182 seconds for the Effluent dataset. N-HiTS demonstrated the second-best performance with 198 seconds, followed by LSTM (542 seconds) and ARIMA (2992 seconds). This experiment underscores the potential of our proposed method to enhance computational time efficiency, making it suitable for adoption in real-time surveillance systems.

**Table 3.**
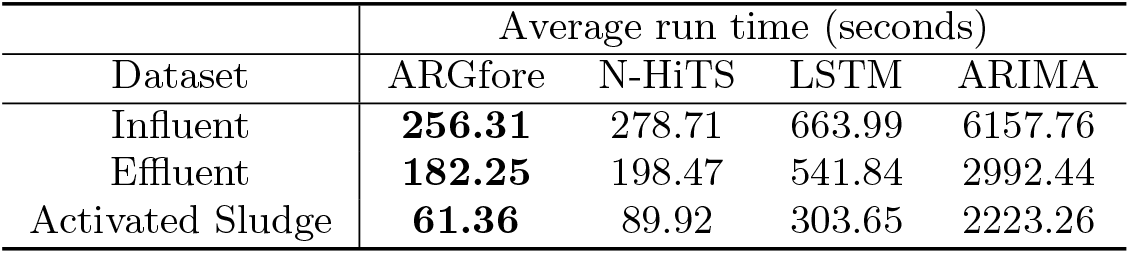
Average run time of ARGfore with the comparison methods for time-series forecasting.

### 3.4 Forecasting performance under varied sampling frequency

To offer researchers guidance for decision-making regarding sampling frequency in predicting future ARG abundances, we explored how the forecasting performance varies under different sampling frequencies. The Effluent and Influent dataset was divided into an 8 : 2 ratio for training and testing the model. Activated Sludge dataset was excluded from this experiment due to its monthly sampling frequency, given that the other datasets were collected twice per week. For the training dataset, MICE imputation was conducted to prepare multiple datasets, each featuring a different regular day interval ranging from 3 to 7 days. MAPE and SMAPE were then measured for the testing dataset, and the experiment was repeated five times. Figure 4 shows a significant increase in prediction error for future ARG abundance when the sampling frequency is reduced to intervals of five and six days.

**Fig. 4.**
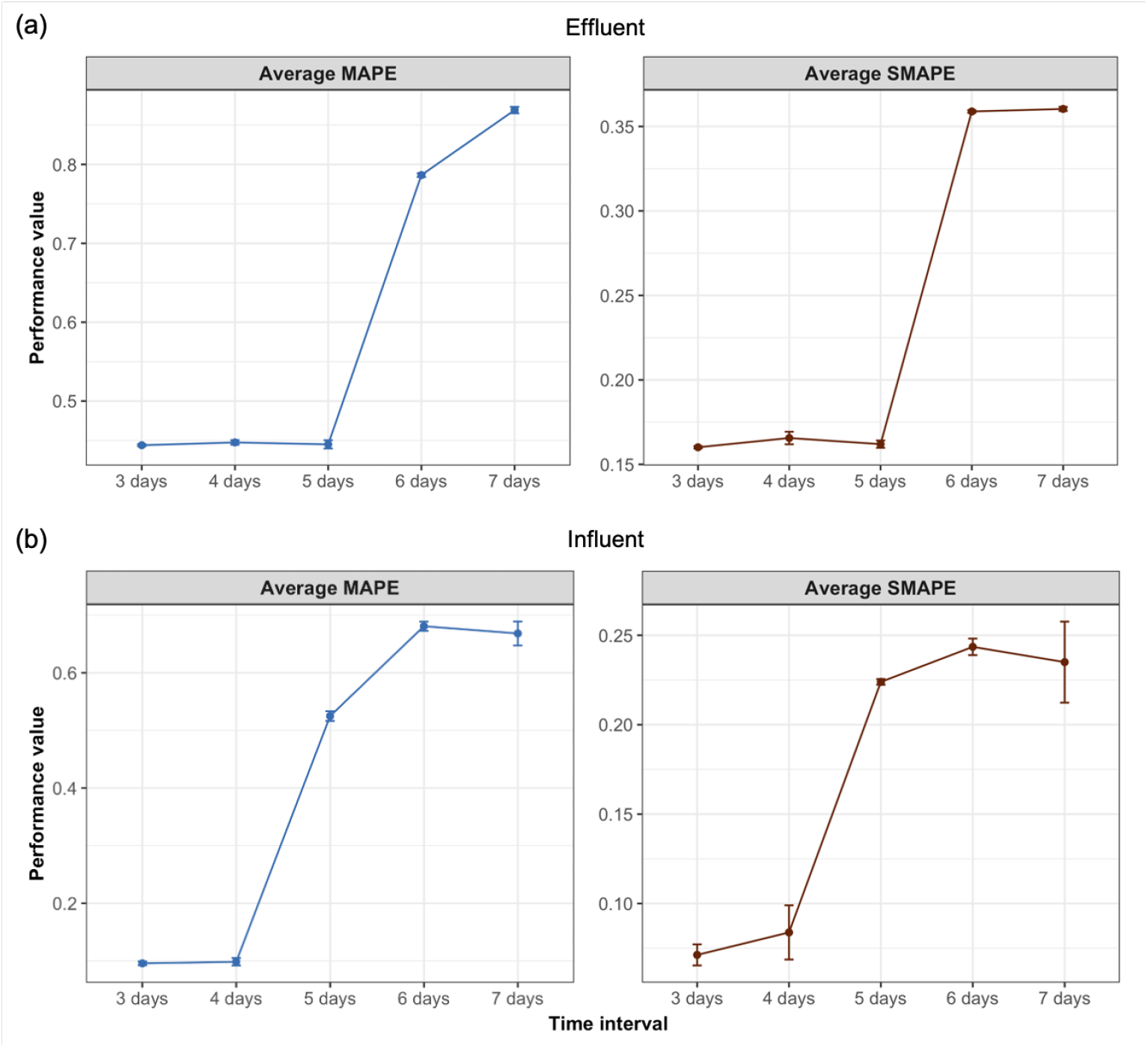
Performance change in time-series forecasting under different wastewater sampling frequency for Effluent and Influent dataset. Each dataset was divided into an 8:2 ratio for training and testing the model, where MICE imputation was conducted to prepare multiple datasets, each featuring a different regular day interval ranging from 3 to 7 days. Prediction error increases significantly when the sampling frequency extends to intervals of five and six days indicating that sampling twice a week provides the more accurate forecasts.

## 4 Discussion and Conclusions

In this paper, we introduced ARGfore, a multivariate time-series forecasting model designed to predict future ARG abundance from metagenomic sequencing data. To the best of our knowledge, this model is the first one to harness the power of deep learning in forecasting ARG abundance using environmental metagenomic data. ARGfore employs a drug-class-based feature extraction module using CNN to capture the inherent relationships between ARGs. These features are then transferred to a forecasting module comprising three MLP-based blocks that decompose the input into trend, seasonality, and residue components. Evaluations using wastewater metagenomic datasets, incorporating different imputation strategies, and comparing run times, demonstrated ARGfore’s improved performance, exhibiting the lowest average MAPE and SMAPE alongside robust computational efficiency. Further, the efficacy of the drug-class-based feature extraction module was tested against various clustering approaches, confirming its ability to support accurate ARG abundance predictions. Additionally, the study explores the impact of different sampling frequencies on ARG abundance forecasting, providing valuable guidance for selecting appropriate sampling intervals for future predictions.

ARGfore has the potential to be seamlessly integrated into any ARG-based surveillance systems, functioning as an early warning system for potential surges in abundance of ARGs of concern. For example, it can be utilized to monitor medically important ARGs listed by World Health Organozation [61]. The capability to predict multiple steps ahead for a diverse set of ARGs positions ARGfore as a valuable tool for proactive interventions and will aid in optimizing treatment processes to counteract the proliferation of antibiotic-resistant microorganisms and their release to the environment. Furthermore, the insights into potential disease outbreaks derived from ARG data can inform policy development, aiming to minimize the environmental dissemination of antibiotic resistance.

One limitation of ARGfore is its implicit assumption that data samples are collected at consistent time intervals. Irregular sampling intervals could pose a challenge for applying ARGfore to forecasting tasks. In response to this limitation, we conducted experiments exploring various imputation strategies for addressing missing data during data preprocessing. Among those, MICE imputation performed the best across all three datasets. The results offer guidance on selecting appropriate imputation techniques to enhance prediction accuracy under varied circumstances. Additionally, we initially selected frequently occurring genes to minimize noise originated from missing data. However, further testing reveals that ARGfore can also be effectively applied on less frequently occurring ARGs. We also examined the impact of sampling frequency on predictive performance, finding that sampling twice per week yielded more accurate forecasts. This framework can be adopted by researchers to determine how frequent or what intervals the sampling should be for monitoring their specific environments [62].

Although ARGfore was developed and trained for forecasting ARG abundance using metagenomic data, its adaptable nature allows potential application to other ARG abundance data collection approaches, such as qPCR. As a method, qPCR is considered to be more sensitive and informative for targeted gene monitoring while metagenomics provides a broader view of the ARGs in microbial communities, enabling exploration of both known and novel genes [63]. ARGfore could potentially be extended to accommodate the qPCR surveillance data where the feature extraction module is not necessary for the single gene target and the forecasting model is trained and developed to predict only one gene’s abundance instead of many. Enhancing ARGfore’s versatility in this way could significantly improve its utility in predicting antibiotic resistance trends across multiple monitoring platforms. Future updates of ARGfore can also include forecasting of antibiotic resistant pathogens.

## List of abbreviations

(ARGs): Antibiotic resistance genes
(AMR): Antimicrobial resistance
(ARIMA): Auto-Regressive Integrated Moving Average
(CARD): Comprehensive Antibiotic Resistance Database
(CNN): Convolutional neural network
(DeepARG-DB): DeepARG database
(FC): Fully connected
(GRU): Gated Recurrent Unit
(LSTM): Long Short-Term Memory
(LOCF): Last Observation Carried Forward
(MAE): Mean absolute error
(MAPE): Mean absolute percentage error
(MGEs): Mobile genetic elements
(MLP): Multi-Layer Perceptron
(MICE): Multivariate imputation by chained equations
(RNN): Recurrent neural networks
(SARIMA): Seasonal ARIMA
(SMAPE): Symmetric mean absolute percentage error
(VARMA): Vector Autoregressive Moving-Average
(WWTP): Wastewater treatment plant

## Declarations

### Ethics approval and consent to participate

Not applicable

### Consent for publication

Not applicable

### Availability of data and materials

Influent and Effluent datasets are available from NCBI (https://www.ncbi.nlm.nih.gov/bioproject) under the accession number PRJNA1083020 and Activated Sludge dataset is available under the accession number PRJNA432264. ARGfore is publicly available at https://github.com/joungmin-choi/ARGfore.

### Competing interests

The authors declare that the research was conducted in the absence of any commercial or financial relationships that could be construed as a potential conflict of interest.

### Funding

This work was supported by the U.S. National Science Foundation (NSF) under Awards #2004751, #2125798, and #2319522, as well as the National Institutes of Health (NIH) grant #1R01AI179686-01A1.

### Authors’ contributions

J.M. Choi designed and implemented the deep learning models, carried out associated experiments and contributed to manuscript writing. M.A. Rumi conceptualized the theory, preprocessed all data, performed statistical experiments and contributed to manuscript writing. A. Pruden and P.J. Vikesland helped supervise the project. L. Zhang supervised the project.

## Acknowledgements

We would like to thank Connor L. Brown and Ayella Maile-Moskowitz for sharing their insights and expertise in biochemistry, which were helpful in designing several experiments.

## Notes

### Competing Interest Statement

The authors have declared no competing interest.

